# Species-specific bleaching trajectories during the 4^th^ global coral bleaching event in northeastern Peninsular Malaysia

**DOI:** 10.1101/2025.10.14.682322

**Authors:** Sebastian Szereday, Kok Lynn Chew, Christian R. Voolstra

## Abstract

During the 4^th^ global coral bleaching event (2023-2025), over 80% of the world’s coral reefs experienced bleaching-level heat stress above 4 °C-weeks (DHW). Inevitably, the large spatial extent of this event substantially impacted coral reefs worldwide, and data are needed to quantify bleaching trajectories and coral mortality. In northeastern Peninsular Malaysia, heat stress accumulated to record levels by June 2024 (DHW=9.6 °C-weeks), resulting in severe mass coral bleaching. Here, we quantified species-specific bleaching and mortality trajectories for 12 abundant reef-building coral species of the genera *Acropora*, *Diploastrea*, *Echinopora*, *Heliopora*, *Montipora*, and *Porites*, by tracking 1850 tagged colonies from September 2023 to October 2024 using a novel Colony Bleaching Response Index (CBRI) that considers scales of discoloration and colony surface extent. In June 2024, 92.9% of all surveyed corals were bleached, resulting in 40.6% mortality by October 2024. However, bleaching trajectories varied across species, whereby the least bleached species in June were not the first to recover nor the least affected by mortality, suggesting inherent species-specific bleaching trajectories and recovery pathways. Mortality of *Porites* cf. *lobata* and *Diploastrea heliopora* was ≤1%, whereas severe mortality was recorded for *Acropora* species (five species, range 46.3-95.4%), *Echinopora* cf. *horrida* (44.7%), and *Montipora* cf. *aequituberculata* (27.8%). Among susceptible species was *Heliopora coerulea*, a species commonly considered heat tolerant. Our findings challenge previous regional studies that concluded a reversal of bleaching hierarchies of species over time, whereby historically ascribed heat tolerant species (i.e., slow-growing massive species) became susceptible and historically susceptible fast-growing species became heat tolerant. Importantly, bleaching severity did not decrease with depth. These data represent the first regional accounts of species-specific coral bleaching mortality in Malaysia, highlighting distinctive ecological bleaching and recovery trajectories of species.

## Introduction

The world’s coral reefs have now experienced four major global coral bleaching events in less than three decades (Hoegh-Guldberg 1999; Hughes et al. 2017, 2018a; Oliver et al. 2018; Eakin et al. 2019; Reimer et al. 2024). While the outcomes of the third global coral bleaching event (2014-2017) are still under investigation (Eakin et al. 2022), unprecedented heat stress unfolded in the Caribbean and Eastern Tropical Pacific during the 2023 boreal summer (Hoegh-Guldberg et al. 2023), leading to severe mass coral bleaching throughout these regions (Neely et al. 2024; Bon et al. 2025; Doherty et al. 2025). Concurrent with the onset of an El Niño Southern Oscillation (ENSO) event in 2023, mass bleaching started to occur in all ocean basins throughout 2024, resulting in the declaration of the 4^th^ global bleaching event (Reimer et al. 2024). In early 2025, the National Oceanic and Atmospheric Administration (NOAA) Coral Reef Watch (CRW) program concluded that between 2023 and early 2025, over 80% of the world’s reefs experienced severe heat stress likely to induce coral bleaching^1^, suggesting that this event is the most widespread global bleaching event in recorded history. This extensive spatial scale requires empirical data to accurately quantify coral bleaching trajectories and associated coral mortality to determine the severity of the 4^th^ global bleaching event.

Malaysian coral reefs are highly diverse and cover widespread areas across the Sunda Shelf and the Sulu Sea marine ecoregions (Huang et al. 2015; Veron et al. 2015). In both regions, economic services derived from coral reef tourism and fisheries are critical for local and regional economies (Sukarno et al. 2015). Unfortunately, both regions have experienced mass bleaching events in the past, concurrent with global bleaching events in 2010 and 2024 (Guest et al. 2012). Between these events, regional-scale bleaching events were also reported during La Niña years (Rosedy et al. 2023; Szereday et al. 2024). Monitoring of these events yielded crucial insights and revealed a reversal of bleaching hierarchies, suggesting that coral taxa historically ascribed as resilient became susceptible to heat stress and vice versa (Guest et al. 2012; Szereday et al. 2024). Secondly, distinctive spatiotemporal patterns in regional bleaching occurrence and severity emerged in the Sunda Shelf region, an extensive shallow continental shelf in Southeast Asia that connects the Malay Peninsula, Borneo, Sumatra, Java, and surrounding islands, whereby reefs in the northeast of Peninsular Malaysia bleached more severely and frequently since 2010 than reefs in the southeast (Szereday et al. 2025a). However, species-specific bleaching susceptibilities and trajectories across these events have not been documented and remain largely unresolved in Malaysia. Understanding the susceptibility of coral species to heat stress across spatial and temporal scales is increasingly important as marine heat stress events increase in severity, scale, and duration (Eakin et al. 2019; Skirving et al. 2019). Regionally, across the east coast of Peninsular Malaysia, the observed north-south gradient in bleaching severity warrants further species-level investigation to identify thermal limits of coral species. Such knowledge could improve regional management and federal policy to better protect coral reefs in Malaysia.

Coral bleaching in response to heat stress is highly dynamic and strikingly complex across all entities of the coral holobiont (Voolstra et al. 2025a). The susceptibility of coral species to heat stress varies across events (Woesik et al. 2011; McClanahan 2017), geographic regions (Sully et al. 2019; McClanahan et al. 2020b), across and within reef sites, across depth, and across micro-environments (Pineda et al. 2013; Baird et al. 2018; Brown et al. 2023). This complexity is further enhanced by differences across genotypes and within a single colony (Drury and Lirman 2021; Voolstra et al. 2025a), complicating in situ assessments of bleaching severity. Within-colony bleaching variability is known for multiple species (Baird and Marshall 2002; van Woesik et al. 2022), and may be underpinned by intra-colony differences in *Symbiodiniaceae* associations (Baker et al. 2008; Hume et al. 2020), and variable microbiome assemblage of the coral-host (Ziegler et al. 2017; Voolstra et al. 2024). To more accurately reflect and capture the complexity of species- and colony-level bleaching outcomes, we devised a categorical metric that accounts for within-colony variability in bleaching response. For example, while a coral may be half bleached (i.e., 50% of the colony surface), the other half is not necessarily healthy or unaffected by heat stress (i.e., it might be pale or dead). Consequently, traditional grouping of colonies into simplified categories, e.g., 0-20% bleached, 21-40% bleached, etc., may obscure bleaching assessments on a colony-level, and further on higher scales when common bleaching indices are averaged across groups (e.g., species, reef sites, depth, etc.). Therefore, this study assessed the outcomes of the most severe heat stress event in northeastern Peninsular Malaysia by investigating within-colony bleaching severity across 12 abundant reef-building coral species. The data provide a previously missing record of species-specific heat stress susceptibilities in Peninsular Malaysia, and ultimately compare bleaching trajectories across reefs and depths to provide a critical update on how distinct hard coral species respond to ocean warming in Peninsular Malaysia.

## Methods

### Reef sites and coral colony tagging

The research was conducted around Pulau Lang Tengah in northeastern Peninsular Malaysia (5°47’43.2’’N, 102°53’39.7’’E Supplementary Figure 1). Species-specific bleaching trajectories of 1,850 tagged colonies from 12 common Indo-Pacific species with varying morphology and life-history strategies were investigated between 1-16 m water depth across five sites (i.e., Batu Bulan - BB, Batu Kucing - BK, House Reef - HR, Karang Nibong - KN, and Tanjung Telunjuk - TT). Surveyed coral colonies were equally distributed across sites, but sample sizes across depth varied across species due to their innate natural distribution along depth gradients (Supplementary Tables 1-2). Despite their proximity to each other (i.e., maximum 500 m distance, Supplementary Figure 1), assessed sites are differentially exposed to wind and wave conditions, resulting in variable diel temperature regimes across sites (Szereday et al. 2024). Furthermore, physical reef health varies across sites and depths, whereby live hard coral cover and density are highest at Batu Bulan and Tanjung Telunjuk, and at shallow depths (Bernard et al. 2023). Coral species were identified based on the distinctive morphological features of the coral skeleton available in online databases (Veron et al. 2019), and cross-referenced with a historical database to verify the species occurrence in this region (Harborne et al. 2000; Veron et al. 2015). However, in lieu of the difficulty and uncertainty associated with assigning species identity based on morphological features assessed underwater (Bridge et al. 2024; Rassmussen et al. 2025), we maintain a non-definitive nomenclature and use the provisional identification term ‘cf.’. In foresight of possible bleaching under the developing El Niño conditions, coral colonies were tagged between September and October 2023 using numbered rubber tags. Each colony was photographed and mapped to ensure redetection during subsequent bleaching surveys in 2024. Importantly, all colonies were visually healthy at the time of tagging, i.e., showed no signs of bleaching, disease, predation and partial mortality. Colonies were spaced at a minimum of 5 m to reduce the assessment of clonal genotypes. Although species-specific sample size was initially standardised across sites, our ability to track the colonies due to loss of tags and coral mortality before bleaching led to varying sample sizes across species (Supplementary Table 1).

### Heat stress metrics

Heat stress during the 2024 coral bleaching event was measured remotely by the National Oceanic and Atmospheric Administration’s (NOAA), Coral Reef Watch product (CRW) version 3.1 (Liu et al., 2014). Satellite measurements were binned to the nearest 5 km^2^ satellite pixel for Pulau Lang Tengah (5°46’30.0"N, 102°52’30.0"E). Sea surface temperature (SST) data were extracted and used to determine heat stress accumulation, commonly described in Degree Heating Weeks (DHW, °C-weeks) (Wellington 2001). Additionally, a refined and locally more representative nDHW metric (DeCarlo 2020; Szereday et al. 2024) was calculated as (equation 1), effectively considering an increase in temperature above the maximum monthly mean (i.e., MMM):

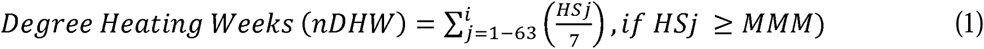

To further describe heat stress levels in situ across sites and depth, ten HOBO ProV2 temperature loggers (Onset Computer Corporation, USA, accuracy level ±0.2°C) were deployed at five sites at 5 and 15 meters on March 17, 2024 (one per site per depth) and set to log at 60-minute intervals. For heat stress determination based on in situ temperature measurements, in situ temperature data were interpolated with the satellite-based maximum monthly mean (i.e., MMM = 29.94 °C). Briefly, the warmest month of the year is routinely in May. All available historical in situ temperature data from all five sites recorded at 8 m were used to calculate the mean monthly temperature in May for the three-year period between 2020 and 2022 (i.e., 30.71 °C). The difference between mean in situ temperature and mean satellite temperature data during this period (i.e., 30.71 – 30.42 °C) was then added to the long-term MMM of 29.94 °C. This interpolation established the in situ MMM of 30.23 °C, which was used for subsequent in situ heat stress calculations based on equation 1 (i.e., nDHW).

### Within-colony bleaching assessment

All reef sites were surveyed between March and October 2024 to track bleaching trajectories of tagged coral colonies throughout the heat stress period. The analysis focused on bleaching survey data collected between June 10 and 21, 2024 (i.e., peak heat stress), and between October 24 and 31 (i.e., four months post-peak heat stress), noting the greater importance of the October data point to determine putative recovery and short-term mortality consequential to heat stress (Claar and Baum 2019). The heat stress response of individual colonies was visually assessed by estimating the percentage of coral colony surface that was bleached. Bleaching was categorized in seven groups: B1 – no bleaching, B2 – pale live, B3 – pale and mild colourful bleaching (i.e., colony visually glowing), B4 – severely pale, B5 – severely pale and severe colourful bleaching, B6 – complete bleaching, B7-dead due to bleaching (Table 1, Figure 2, Supplementary Figure 2). The extent of bleaching per category was noted for each colony in steps of 5%, whereby the sum of all observations added up to 100% (i.e., the entirety of the colony). This percentage-based assessment ensured that within-colony bleaching severity is calculated based on the percentage of bleached colony surface tissue within each category, thus explicitly considering bleaching heterogeneity of colonies instead of compromising such detail into simplified groups. For example, a colony-level bleaching could be 50% bleached, 25% severely pale, and 25% healthy, or 50% bleached, 40% healthy, and 10% dead, etc. These observations were then weighted based on severity by adapting the common Bleaching Response Index (BRI) (McClanahan et al. 2007; Guest et al. 2012; Pratchett et al. 2013; Szereday et al. 2024) to fit colony-level data to calculate the Colony Bleaching Response Index (equation 2):

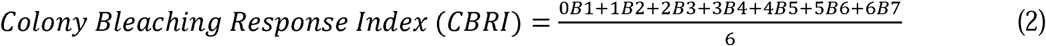

Thus, the CBRI Index is a categorical, normalized, and weighted measure on a 0-100 scale, whereby higher values indicate more severe and complete bleaching across the colony. The CBRI was calculated for all colonies for both bleaching data points, June 2024 and October 2024. The species-specific CBRI values were further averaged by species, site, and depth to compare bleaching severity across and within reef-scales.

**Table 1.**
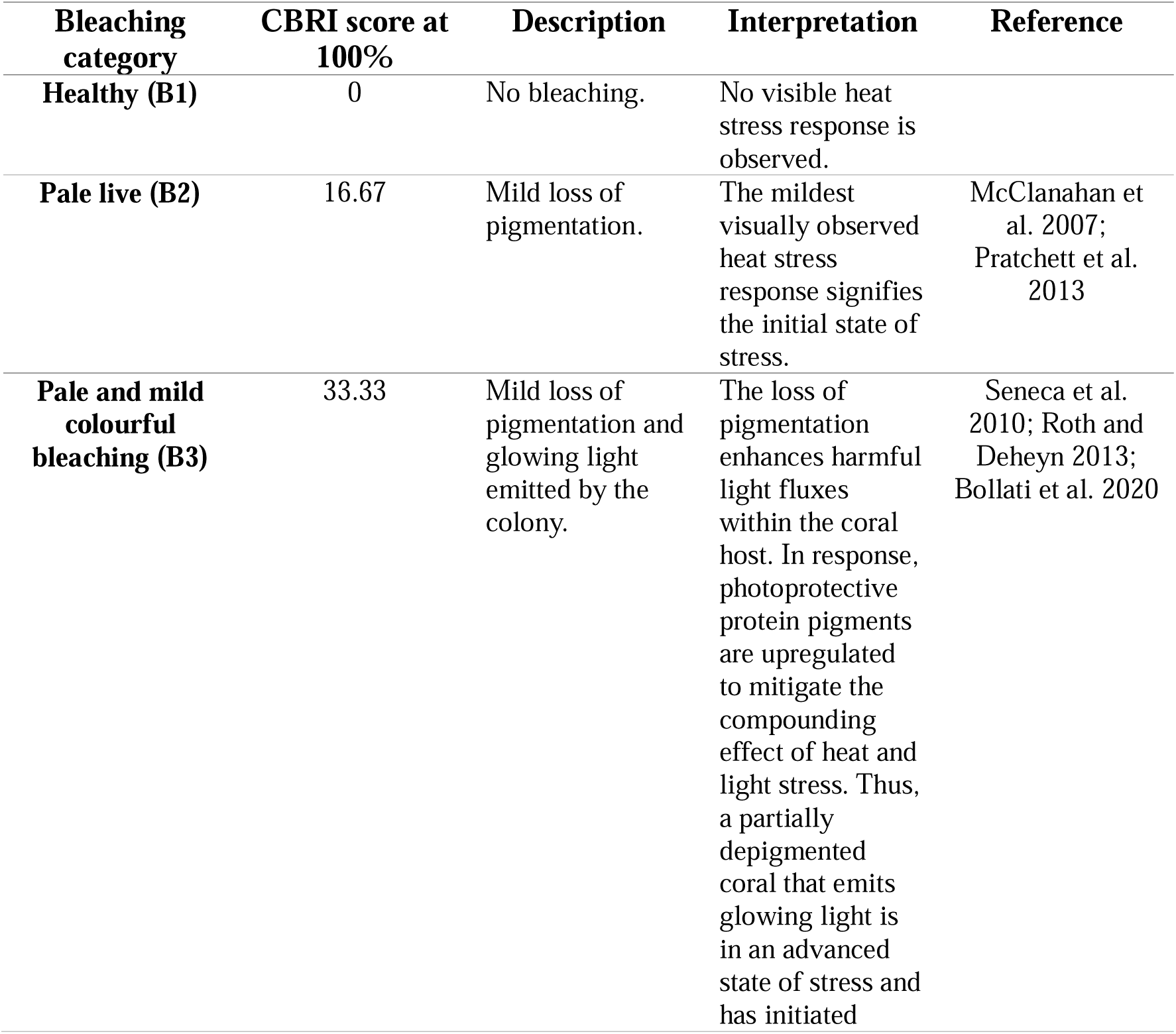
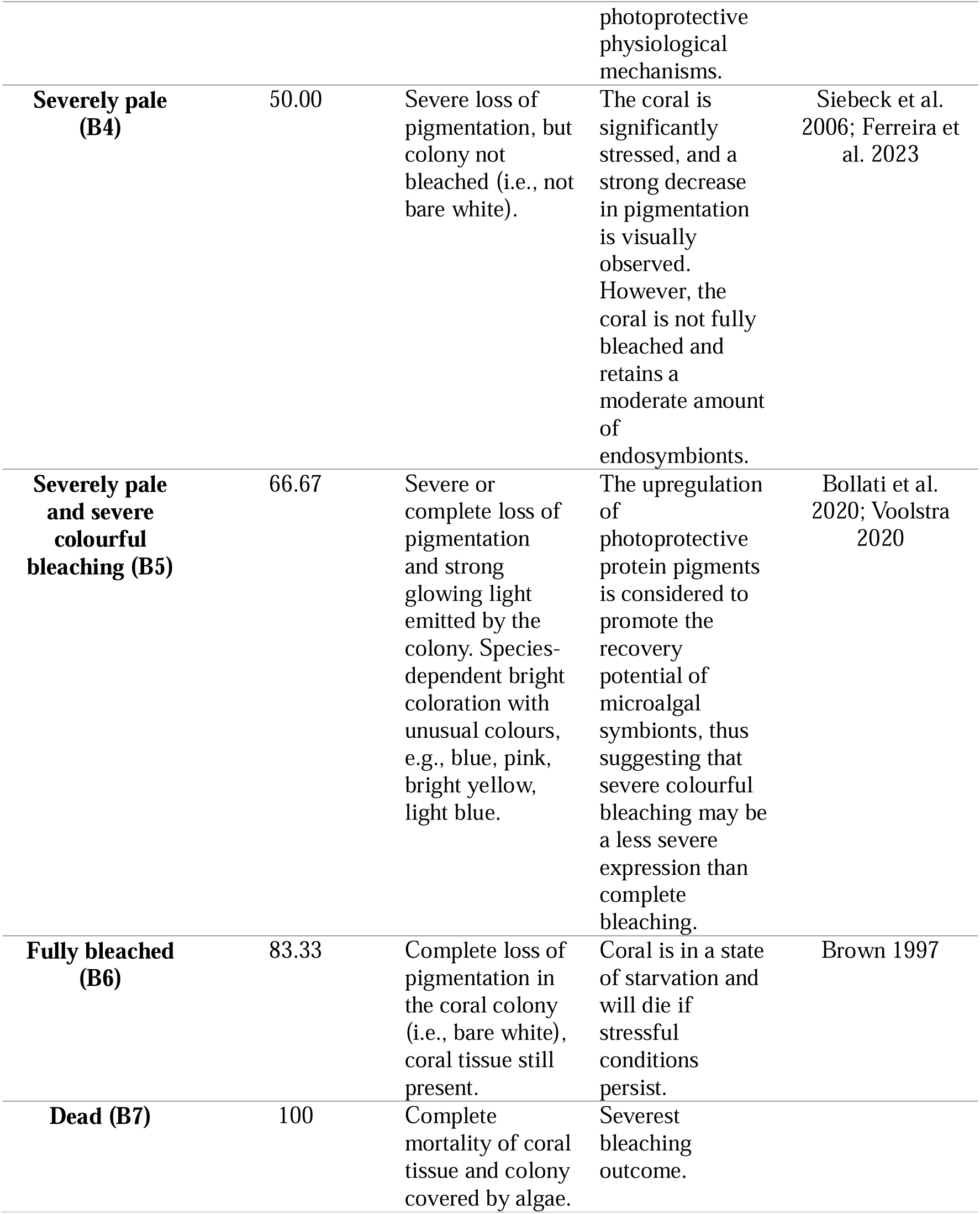
Description and interpretation of coral bleaching categories at colony-level.

### Statistical analysis

All statistical analyses were conducted at the species level, for both survey time points (i.e., CBRI in June and October), at a significance level of p ≤ 0.05. To assess bleaching severity across depth, coral colonies were divided into the following depth categories: Shallow (1-5.4 m), Intermediate (5.5-10 m), and Deep (> 10 m). A general linear model (GLM) with a binomial distribution was used to examine predictors of bleaching. Specifically, we examined how species-specific bleaching risk varied by site and depth by estimating marginal means from the fitted GLMs to provide group-level comparisons (i.e., predicted bleaching probability). To account for within-colony variability, CBRI scores were categorized into seven groups by severity: B1 (CBRI=0), B2 (CBRI= 0-16.7), B3 (16.8-33.3) B4 (33.4-50), B5 (50.10-66.7), B6 (66.8-83.3), B7 (>83.3), and a logit-link function was incorporated into the model to transform the dependent variable (i.e., CBRI June and October) into binomial distribution probabilities for GLM modelling (Nelder and Wedderburn, 1972). GLM analyses were performed in R Studio version 4.3.2 (R Core Team, 2023) using the packages ‘*modEVA*’ (Barbosa et al. 2014), ‘*car*’ (Fox and Weisberg, 2018), and ‘*emmeans*’ (Lenth 2025). In lieu of non-normal distribution and outliers, mean CBRI differences across site and depth were tested with Kruskal-Wallis tests followed by Dunn’s pairwise multiple comparisons with Holm’s correction of p-values. Finally, time-series heat stress data (i.e., nDHW) specific to each site and depth were examined using a Kruskal-Wallis test followed by Dunn’s test for multiple comparisons.

## Results

### Unprecedented, record-breaking heat stress in 2024

The 2024 heat stress period started in April and reached maximum levels between June 18, 2024 (DHW=9.6 °C-weeks) and June 21, 2024 (nDHW=10.9 °C-weeks) (Figure 2). Previous record levels (DHW=5.2 °C-weeks and nDHW=8.6 °C-weeks) measured in 2010 were exceeded by 4.4 °C-weeks (DHW) and 3.3 °C-weeks (nDHW), respectively (Figure 1, Supplementary Figure 3). In situ heat stress levels were significantly greater at shallow depth at 5 m (i.e., mean nDHW 11.5 °C-weeks) compared to 15 m (mean nDHW 9.7 °C-weeks) across all sites (Kruskal-Wallis H=67.57, df=8, p<0.001) (Figure 2). Importantly, heat stress was not significantly different across sites when comparing equal depths (i.e., shallow vs. shallow, deep vs. deep), except for Tanjung Telunjuk at 15 m (i.e., TT Deep), where heat stress was significantly lower (i.e., 9.1 °C-weeks) compared to all other sites, regardless of depth (Figure 2, Supplementary Table 3). Diel temperature profiles across sites and depth are accessible in Supplementary Figure 4.

**Figure 1.**
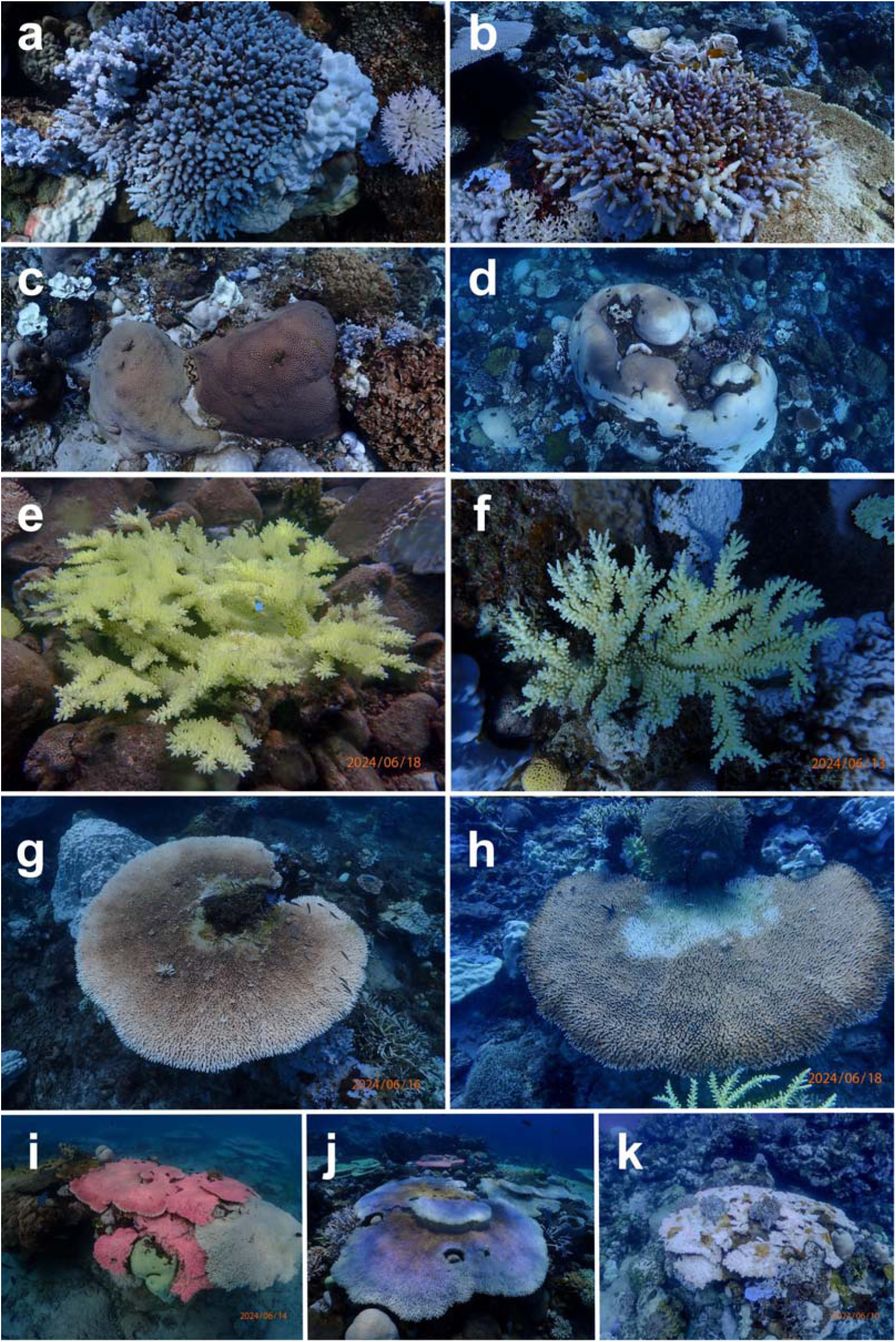
Example of coral bleaching categories. (a-b) *Acropora* cf. *gemmifera* colonies showing differential bleaching expressions: (a) severely pale and severe colourful bleaching vs. (b) severely pale, severe colourful bleaching, and fully bleached branches. (c) A pale *Diploastrea heliopora* colony (left colony) next to a healthy colony (right colony). (d) Partial bleaching of a *Diploastrea heliopora* colony. (e-f) Differential colourful bleaching example observed for *Acropora* cf. *florida* colonies: (e) pale and mild colourful bleaching, and (f) severely pale and severe colourful bleaching. (g-h) within-colony bleaching variability seen for *Acropora* cf. *cytherea*: (g) inner part of the table is pale, becoming severely pale towards the outside, where the edge is fully bleached. (h) The fully pigmented edge of the colony becomes severely pale and bleached towards the inside of the colony. (i-j) show differential colourful bleaching among table coral species: (i) mild colourful bleaching of *Acropora* cf. *spicifera* (bright red-pink colony) and (j) severe colourful bleaching of *A. cytherea* glowing in bright blue. Image (k) shows a fully bleached *A. spicifera* colony with partial mortality from heat stress exposure. Images by Chew Kok Lynn and Sebastian Szereday.

**Fig. 2.**
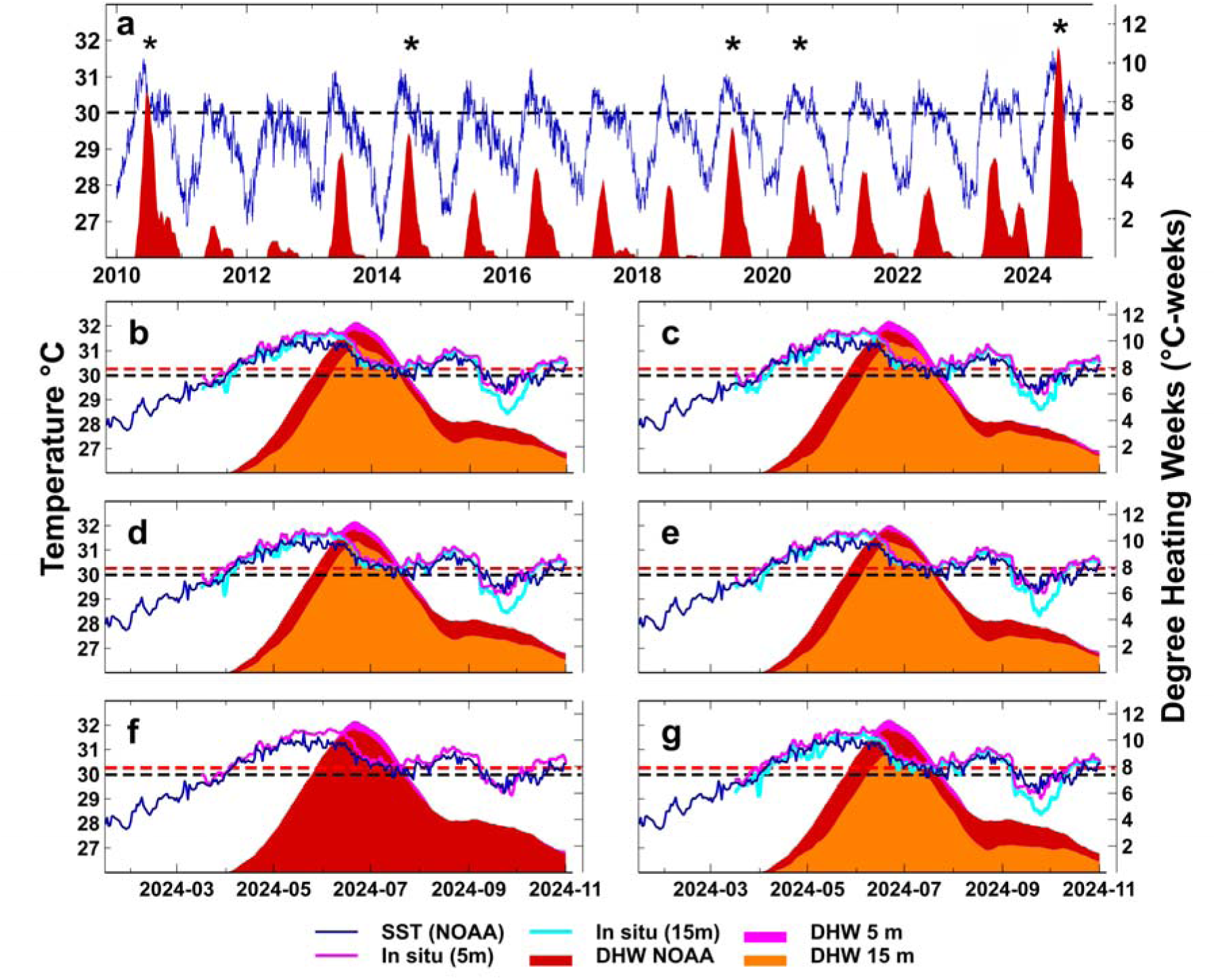
Sea surface temperature (SST) and degree heating weeks (nDHW). The blue line shows the satellite-based nightly sea surface temperatures (SST °C) between 1 January 2010 and 31 October 2024, around Pulau Lang Tengah (5°46’30.0"N, 102°52’30.0"E), recorded by the National Oceanic and Atmospheric Association (NOAA), Coral Reef Watch (CRW) product, version 3.1. The black-dotted line shows the maximum monthly mean (MMM) temperature based on NOAA CRW satellite data (i.e., 29.94 °C). Heat stress based on satellite SST data (i.e., DHW NOAA) is shown in dark red to compare the severity of heat stress events since 2010. The asterisks in panel (a) highlight heat stress events that resulted in coral bleaching (identified from available literature and author observations). Panels (b-g) show site-specific temperature and heat stress data, where the mean daily 24h in situ sea temperature is shown for measurements at 5 m (cyan line) and 15 m (blue line) water depths, respectively. The red-dotted line highlights the MMM (i.e., 30.23 °C) based on in situ temperature data from five sites around Pulau Lang Tengah recorded between 2020 and 2022. nDHW based on in situ temperature data is shown for both depths (DHW 5m and DHW 15m). (b) – island average; (c) – Batu Bulan; (d) – Batu Kucing; (e) – House Reef; (f) – Karang Nibong (deep data missing due to logger malfunction); (g) – Tanjung Telunjuk.

### Distinct species-specific bleaching trajectories

The temporal tracking of coral colonies across depths and sites revealed differential bleaching trajectories and susceptibilities of coral species during the 2024 heat stress period. Bleaching was first noted in early May 2024 and reached severe levels by early June 2024. During surveys conducted between June 10 and 21, at peak heat stress levels, over 92.9% of colonies were bleached at variable levels (Table 2, Figure 3). At this point, *Heliopora coerulea* colonies were experiencing the most and severest bleaching of all species (i.e., 86.0% of colonies fully bleached, mean CBRI = 80.1), followed by *Echinopora* cf. *horrida* (i.e., 17.1% of colonies fully bleached, CBRI = 57.5), and *Montipora* cf. *aequituberculata* (i.e., 20.8% of colonies fully bleached, CBRI = 53.5). In contrast, only 1.8%, 1.7%, and 0.1% of colonies of the three most tolerant species in June, i.e., *Acropora* cf. *cytherea* (CBRI=28.6), *Acropora* cf. *florida* (CBRI=30.9), and *Echinopora* cf. *pacificus* (CBRI=31.7), were fully bleached. However, based on the October data point, we identified lag responses of multiple species to heat stress (Figure 4-5). Specifically, in October 2024, mortality rates of the two most tolerant species in June, i.e., *A. cytherea* and *A. florida*, were very high at 75.9% and 64.6%, respectively, while complete recovery was very low (i.e., 4.6% and 3.4%, respectively) (Table 2, Figure 4). A similar pattern was identified for all other *Acropora* species, as mortality rates of *Acropora* species were the highest among all species, followed by *E. horrida* (44.7%). Particularly, *Acopora* cf. *spicifera* experienced extremely severe levels of mortality at 95.4%. In contrast, slow growing massive species *Diploastrea heliopora* and *Porites* cf. *lobata* bleached moderately in June (CBRI = 33.5 and 44.8, respectively), whereas mortality rates of these species were the lowest overall (i.e., ≤1%). Importantly, for *D. heliopora* and *P. lobata*, 55.8% and 54.9% of colonies, respectively, were fully healthy four months after peak heat stress in October 2024 (Table 2, Figures 3-4). Other species with low mortality included *Porites* cf. *rus* (10.6%), *Echinopora* cf. *pacificus* (13.6%), and *Heliopora coerulea* (13.6%). However, owing to the widespread prevalence of partial and full colony mortality (i.e., 78.3% of all colonies, Table 2), recovery rates were generally low (except for *D. heliopora* and *P. lobata*), ranging from 0% (i.e., *A. spicifera*) to 21.3% (*M. aequituberculata*) (Table 2, Figure 3-4). Taken together, these data show distinctive ecological bleaching trajectories of species, whereby the last species to bleach severely were not the first species to recover nor the least affected by mortality (Figure 3-5).

**Table 2.**
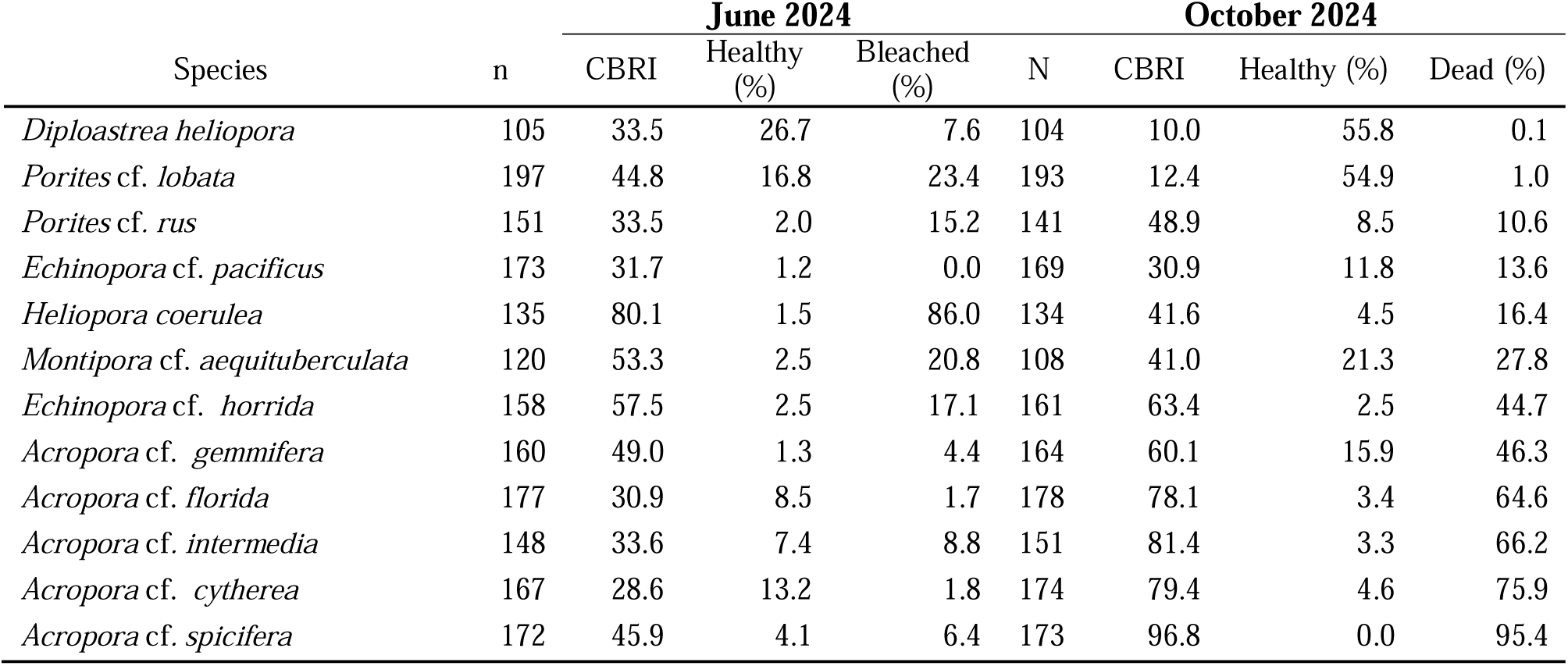
Bleaching and mortality of surveyed colonies in June (n) and October (N) 2024. Percentage of colonies observed as healthy (i.e., no bleaching), bleached (i.e., >90% of colony surface bleached), and dead (>90% of colony surface dead) is shown for each species during peak heat stress in June 2024 and four months after heat stress in October 2024. Species are listed from the lowest to highest bleaching mortality based on the October 2024 data.

**Fig. 3.**
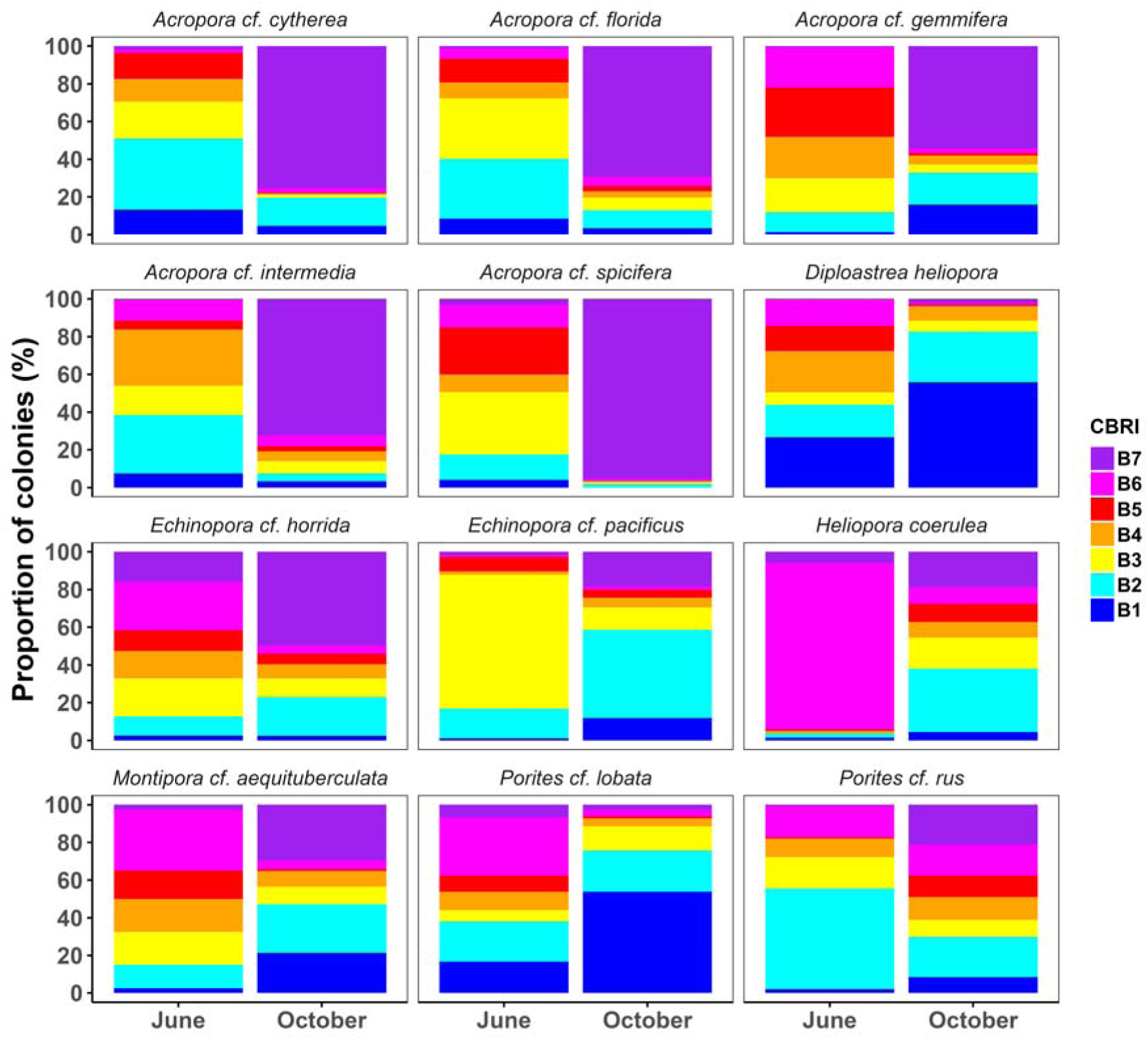
Coral bleaching severity and recovery. The severity of bleaching on colony-level is expressed by the Colony Bleaching Response Index (CBRI) based on data recorded in June and October 2024. Higher CBRI scores indicate more severe bleaching. CBRI categories: B1 – no bleaching, B2 ≤ 16.7, B3 =16.8-33.3, B4 = 33.4-50, B5 = 50.1-66.7, B6 = 66.8-83.3, B7 - CBRI > 83.3. Colour scale based on NOAA’s CRW heat stress alert colour scheme.

**Fig. 4.**
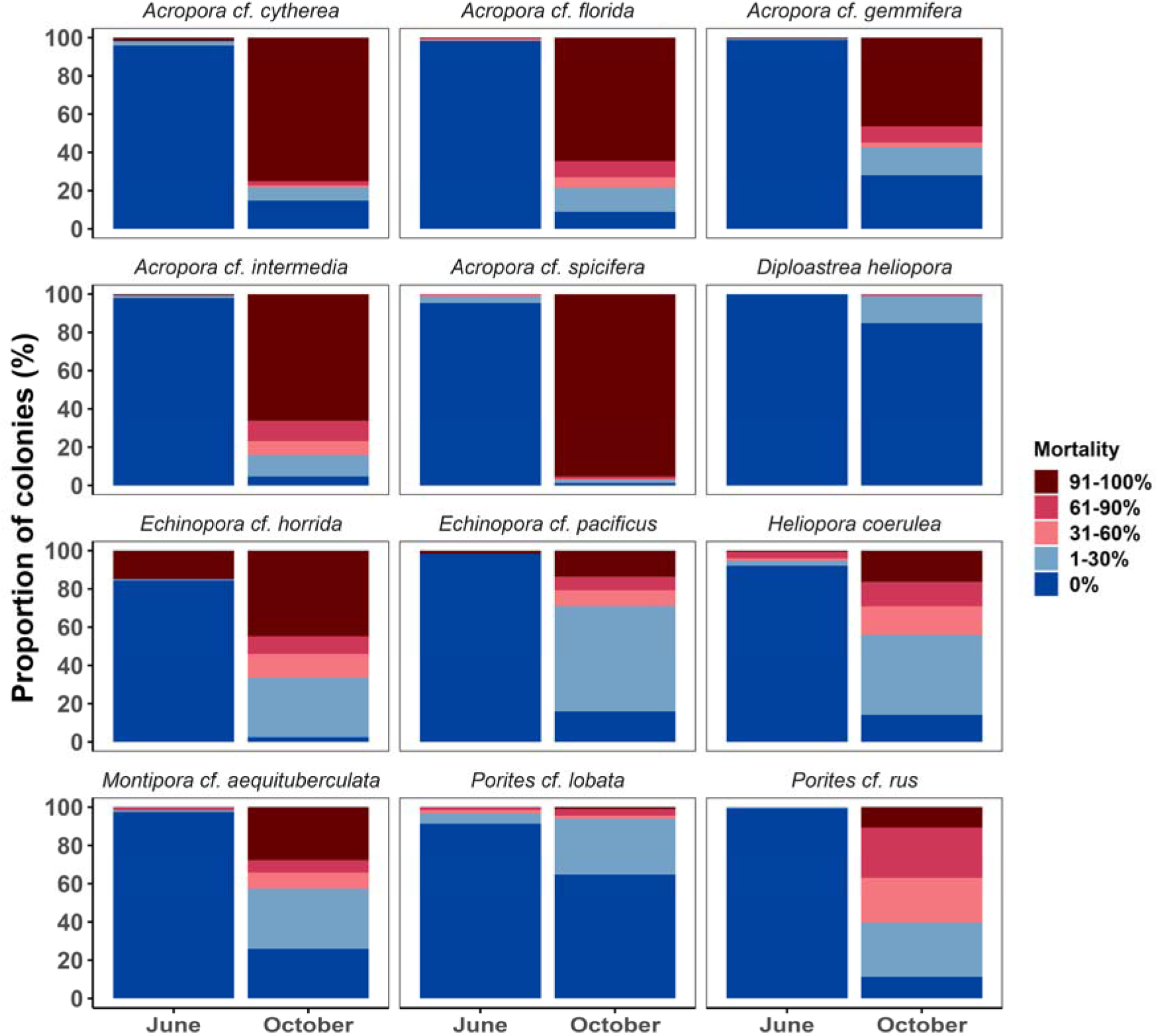
Percentage of healthy, partially, and fully dead coral colonies during peak heat stress in June 2024 and four months later in October 2024.

**Fig. 5.**
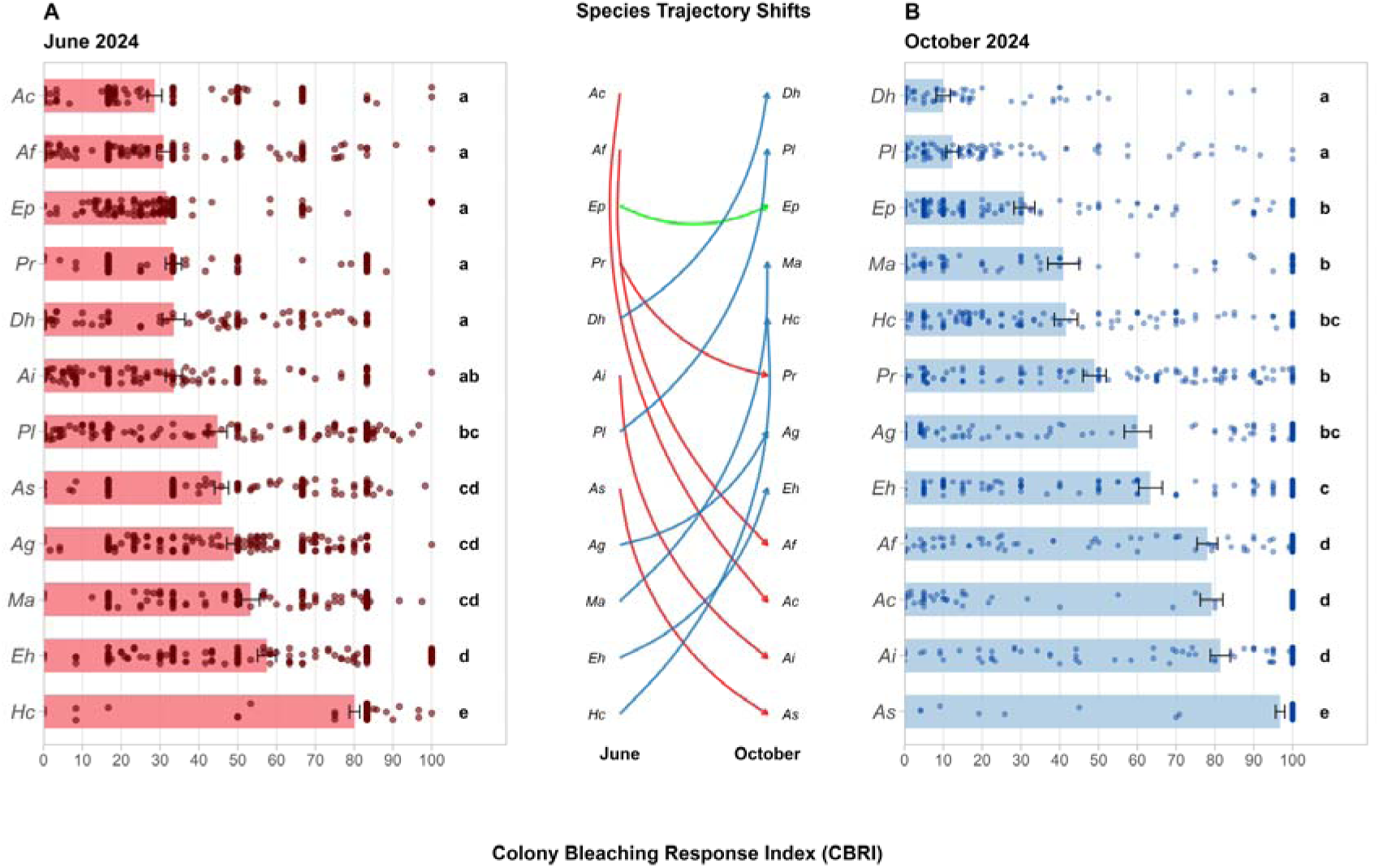
Changes in bleaching trajectory among species across survey periods. Coral bleaching severity is expressed by the Colony Bleaching Response Index (CBRI). Dots represent individual data points of colonies and emphasize within species variability and data distribution. Error bars signify the standard measurement error. Species are plotted from least (top) to worst bleached (bottom) for each survey occasion to emphasize the distinct bleaching trajectories of species. Letter annotations represent results of pairwise Dunn’s Tests conducted at significance levels of p ≤0.05. Species abbreviations: Ac – *Acropora* cf. *cytherea*; Af – *Acropora* cf. *florida*; Ag-*Acropora* cf. *gemmifera*; Ai – *Acropora* cf. *intermedia*; As – *Acropora* cf. *spicifera*; Dh – *Diploastrea heliopora*; Eh – *Echinopora* cf. *horrida*; Ep – *Echinopora* cf. *pacificus*; Hc – *Heliopora coerulea*; Ma – *Montipora* cf. *aequituberculata*; Pl – *Porites* cf. *lobata*; Pr – *Porites* cf. *rus*.

### Limited depth refuge during severe heat stress

At record-breaking heat stress levels, the estimated marginal mean probability of bleaching was predominantly consistent across sites and depths across the examined species (Supplementary Figures 5-7). Nonetheless, notable exceptions existed. For example, bleaching mortality was significantly lower for *Acropora* cf. *gemmifera* (Kruskal-Wallis H=14.48, df=2, p<0.001) and *E. horrida* (Kruskal-Wallis H=11.92, df=2, p=0.0026) at greater depth, while the opposite pattern was found for *P. lobata*, as more colonies bleached in June and died by October at greater depth (Kruskal-Wallis H=36.00, df=2, p<0.001) (Supplementary Figure 5 and Supplementary Table 4). Significant mean October CBRI differences across sites were found for *A. cytherea* (lower CBRI at Karang Nibong), *A. gemmifera* (lower CBRI at Karang Nibong and Tanjung Telunjuk), and *A. florida*, *E. horrida,* and *M. aequituberculata* (higher CBRI at House Reef). Taken together, bleaching differences across depths and sites were limited and suggest little refuge from severe heat stress and bleaching across and within reef-scales (Supplementary Figures 5-7, Supplementary Tables 4-5).

## Discussion

The 2024 heat stress event is the most severe marine heatwave on record for northeastern Peninsular Malaysia and resulted in substantial region-wide mortality across coral species (Szereday et al. 2025a). Our temporal analysis of 1,850 coral colonies revealed very high levels of mortality, with 78.3% of colonies suffering at least partial mortality and 40.7% of colonies dying off entirely. In addition, only 14.7% of corals recovered fully by October or remained unbleached throughout the event. Furthermore, mortality may have increased post monitoring and surveys likely did not capture the full extent of long-term mortality following severe bleaching. Although bleaching mortality data from the previous record heat stress event in 2010 are unavailable, these data suggest that the 2024 event was one of the most severe bleaching events in this region. The extensive mortality of fast-growing species (i.e., *Acropora*) will arguably result in lasting ecological changes and homogenization of coral communities, leading to post-bleaching phase shifts (Bruno et al. 2009; Hughes et al. 2018b; Cannon et al. 2021). On the one hand, high levels of mortality resulted in almost complete elimination of some *Acropora* species and *E. horrida,* with consequences for the prospect of natural recovery via sexual reproduction (Hughes et al. 2019; Leinbach et al. 2021). On the other hand, large amounts of dead biomass have negative legacy effects on recovery rates of all species regardless of bleaching trajectories (Kopecky et al. 2024), and may permanently halt or slow coral reef recovery in this region (Bernard et al. 2023).

In comparison to the most recent bleaching event in 2019 (Szereday et al. 2024), full colony mortality rates increased from 1.1% in 2019 to 40.7% in 2024, in concordance with the higher heat stress loading in 2024 (i.e., 2019 nDHW 6.8 °C-weeks vs 10.9 °C-weeks in 2024). Accordingly, coral bleaching trajectories of species greatly differed under severe heat stress in 2024 compared to moderate heat stress in 2019. Specifically, in 2019, branching *Acropora* and laminar *Montipora* species were largely tolerant to prevailing heat stress levels, whereby these taxa were severely affected in 2024. In contrast, while several species of encrusting and massive *Porites* were bleached substantially in 2019, these species were more tolerant (i.e., less bleaching and mortality) and more resilient (i.e., higher recovery) in 2024 compared to other species. This pattern was observed across the entire Terengganu Island archipelago in northeastern Peninsular Malaysia in 2024 (Szereday et al. 2025a), but not for southeastern Peninsular Malaysia, where heat stress was moderate (i.e., 5.4 °C-weeks) and did not exceed the putative mortality threshold of DHW > 8 °C-weeks (Eakin et al. 2010). Therefore, the putative reversal of bleaching hierarchies since 1998 and 2010 suggested for south and northeastern Peninsular Malaysia (Guest et al. 2012; Szereday et al. 2024) may only be observed at moderate heat stress levels, while these contrasting bleaching response patterns of species suggest differential physiological responses and coping mechanisms of species across heat stress gradients. Alternatively, prior exposure to heat stress may determine future responses to thermal stress (legacy effects), which may produce complex bleaching susceptibility dynamics over time, as suggested earlier (Evensen et al. 2022). Irrespective of this, slow growing species such as *P. lobata* and *D. heliopora* responded early to heat stress and bleached first. Importantly, however, the early onset of bleaching may not posit low thermal tolerance per se but rather reflects a resilient response pattern in lieu of the magnitude and duration of heat stress. For instance, early bleaching may enable temporal symbiont shuffling beneficial for recovery and survival (Cunning et al. 2015; Silverstein et al. 2015), while reducing excess symbiont density may decrease heat stress susceptibility under stressful conditions (Cunning and Baker 2013). Counter to that, fast growing species such as *A. cytherea* and *A. florida* were the last to bleach severely, yet suffered severe mortality rates across all reef environments. This reflects a more static pattern and indicates a putative breaking point after which the corals cannot further attune (Voolstra et al. 2021b). Therefore, species demonstrated distinctive bleaching trajectories and the reversal of bleaching hierarchies compared to 2019 may be explained by the magnitude of prevailing heat stress and the subsequent physiological response patterns, besides putative legacy effects discussed above. Globally, coral taxa are thought to have become more heat tolerant (McClanahan et al. 2020a; Lachs et al. 2024; Howells et al. 2025). However, and importantly, our observational data suggest that such (selection for) thermal tolerance increases ultimately may not translate into higher survival rates if the magnitude and duration of heat stress continue to break records (Frieler et al. 2013). For other species, such as *H. coerulea* and *E. horrida,* bleaching patterns were similar to those in 2019, with a high level of early bleaching and subsequent moderate to high mortality contingent with heat stress severity (i.e., higher mortality in 2024). In the case of *H. coerulea*, these data substantiate that *H. coerulea* is locally susceptible to heat stress, contradicting studies that have identified *H. coerulea* as superiorly heat tolerant (Paulay 1999; Kayanne et al. 2002; Schuhmacher et al. 2005; Phongsuwan and Chansang 2012; Raymundo et al. 2019). Ultimately, for most species, moderate heat stress events such as 2019 are likely not representative nor accurate indicators of species-specific heat stress tolerance and resilience, suggesting that bleaching susceptibilities should not be inferred based on moderate heat stress events.

Depth and other micro-environmental differences have been correlated with differential bleaching of species (Muir et al. 2017; Baird et al. 2018; Brown et al. 2023). Previously in 2019, significant differences across these reef sites and depths were found for numerous *Acropora* species and *H. coerulea* (Szereday et al. 2024; Henry et al. 2025). However, despite lower heat stress at increasing depths (Figure 2, Supplementary Table 3), no discernible differences were found in 2024 for these species (except *A. gemmifera*). Contrarily, bleaching severity of *P. lobata* was lower at shallow depth, consistent with observations from 2019 (Szereday et al. 2024), and further corresponding with observations from other studies (Baird et al. 2018; Crosbie et al. 2019). Ultimately, depth offered limited refuge from heat stress and mitigated bleaching only for *A. gemmifera* and *E. horrida*, while no consistent site-specific pattern was found, which partially opposes previous findings across these reefs during moderate heat stress (Szereday et al. 2024). This can partly be explained by equal heat stress loads across sites; however, the mechanisms underpinning differences across and within species, such as symbiotic associations (LaJeunesse et al. 2018; Santoro et al. 2025), remain unresolved and require urgent investigations.

Owing to the demise of coral reefs worldwide, great interest persists in identifying and selecting naturally occurring heat tolerant and resilient corals for conservation and restoration efforts (Morikawa and Palumbi 2019; Caruso et al. 2021; Roper et al.; Szereday et al. 2025b). However, the distinct bleaching trajectories of species reported here strongly suggest that heat stress tolerance and resilience should not be inferred from a single data point to avoid obscuring coral thermal tolerance assessments (Claar and Baum 2019), which should also be considered during thermal acute assay screening (Szereday 2025b). For example, although there was no significant difference in mean bleaching severity for *A. cytherea*, *A. florida*, and *A. intermedia* compared to *D. heliopora* in June, highly significant differences emerged by October, clearly highlighting distinct trajectories. Thus, due to the response differences of species, multiple data points are ideally required that distinguish bleaching susceptible and tolerant colonies to correctly predict long-term bleaching trajectories. Crucially, colony-level assessments should move beyond binary bleaching metrics (i.e., bleached vs unbleached) and simplified categories (i.e., 20%, 40%, 60% bleached). Instead, the dynamic stages of bleaching (i.e., pale, severe pale, colourful bleaching, etc.) should be considered using integrated metrics such as the Colony Bleaching Response Index (Voolstra et al. 2025b). This adapted bleaching metric captures the complexity of species- and colony-level bleaching outcomes and provides an improved methodology to assess coral thermal tolerance in situ.

The outcomes of the 2024 coral bleaching event suggest a bleak outlook for corals in northeastern Peninsular Malaysia. Beyond heat stress, reefs are threatened by a suite of stressors, and these compounding effects may further erode the adaptive capacity of corals and reduce overall reef resilience. It is worthwhile to note that the temporal tracking of coral colonies on species levels provides critical direction for conservation, as numerous heat tolerant colonies, i.e., colonies that did not bleach or only bleached at minor levels, were identified for all species except *A. spicifera* and *E. horrida*. This opens a window of opportunity to gain insight into the mechanistic underpinnings of thermal resilience, besides such colonies providing an opportunistic starting point for extending the natural adaptive capacity of corals to heat stress (Voolstra et al. 2021a). At this conjuncture, it is critical to resolve what enables these colonies to resist, tolerate, and recover from heat stress to develop strategies and active interventions to conserve species in the wake of unhalted climate change.

## Supporting information

Electronic Supplementary Material (ESM) accompanying the article

## Acknowledgement

Research was conducted under permit number Prk.ML.630-7Jld.14 (1) issued by the Department of Fisheries (DoF) Malaysia (Jabatan Perikanan Malaysia), and permit number EPU 40/200/19/3717 (11), issued by the Ministry of Economy (Kementerian Ekonomi) Malaysia. Our gratitude is extended to Summer Bay Resort Lang Tengah Island, and the entire dive centre crew for their continued support of field research activities. Thank you to our research assistants, Febrianne Sukiato, Kiu Yee Tong, and Dharshinie Mano, for assisting with field work, and to Natasha Zulaikha Zahirudin, for helping with project administration and image preparation.

## Author Contributions

SS conceived and designed the study, conducted the fieldwork, analysed the data, and wrote the original draft; CKL conducted the fieldwork, assisted with data processing and interpretation, and edited the original draft; CRV supported scientific framework development, data interpretation, and edited the original draft. All authors reviewed and edited the final draft and granted approval to submit for publication.

## Data and code availability

All codes and data required to replicate the study are available online at https://github.com/Coralku/bleaching_2024

