## Supplementary material for "Species-specific bleaching trajectories during the 4^th^ global coral bleaching event in northeastern Peninsular Malaysia": Electronic Supplementary Material (ESM) accompanying the article

[1]Coralku Solutions, Non-profit organization for coral reef research and restoration, Kuala Lumpur, Malaysia.

[2]Department of Biology, University of Konstanz, Konstanz, Germany.

*** Correspondence:**
Sebastian Szereday



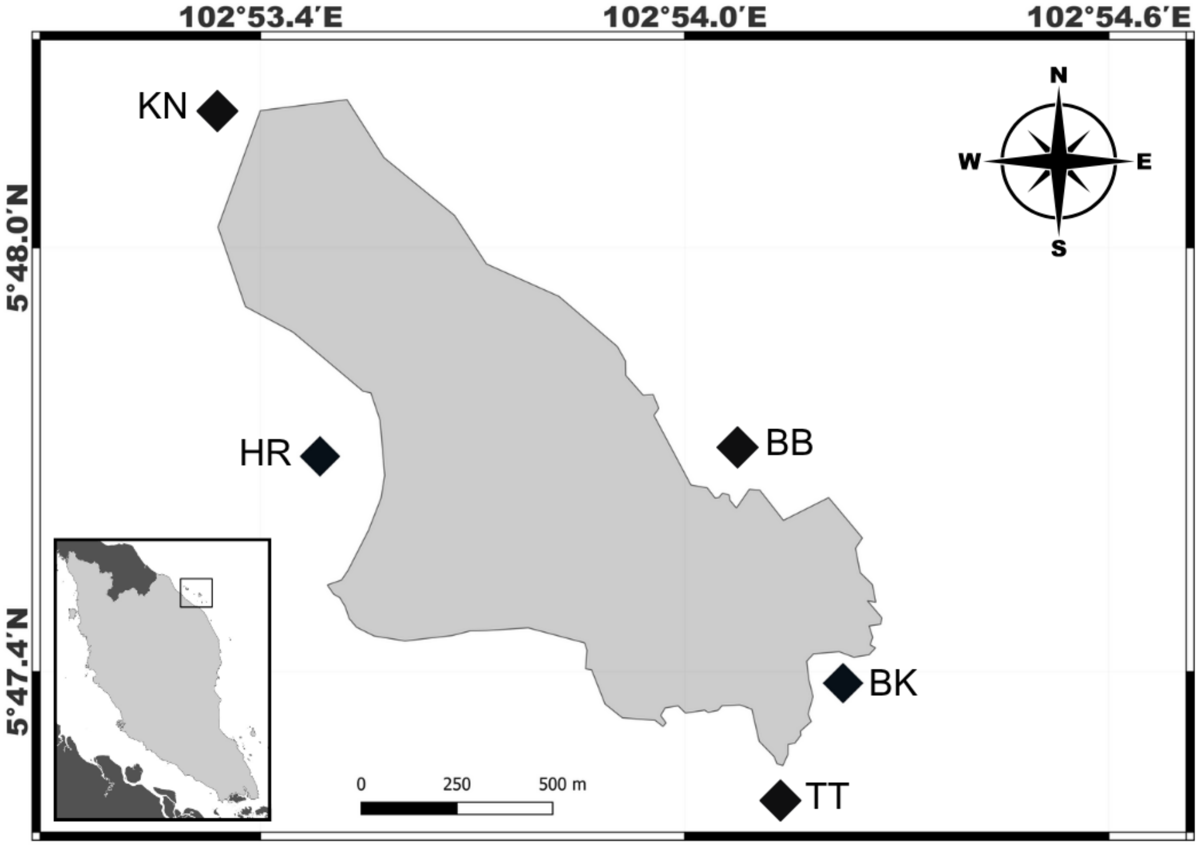


**Supplementary Figure 1. Geographic location of study sites around Pulau Lang Tengah (5°47' 43.2"N, 102°53'39.7"E) along the northeastern coast of Peninsular Malaysia (zoomed out panel).** Site abbreviations BB – Batu Bulan, BK – Batu Kucing, TT – Tanjung Telunjuk, HR – House Reef, KN – Karang Nibong.

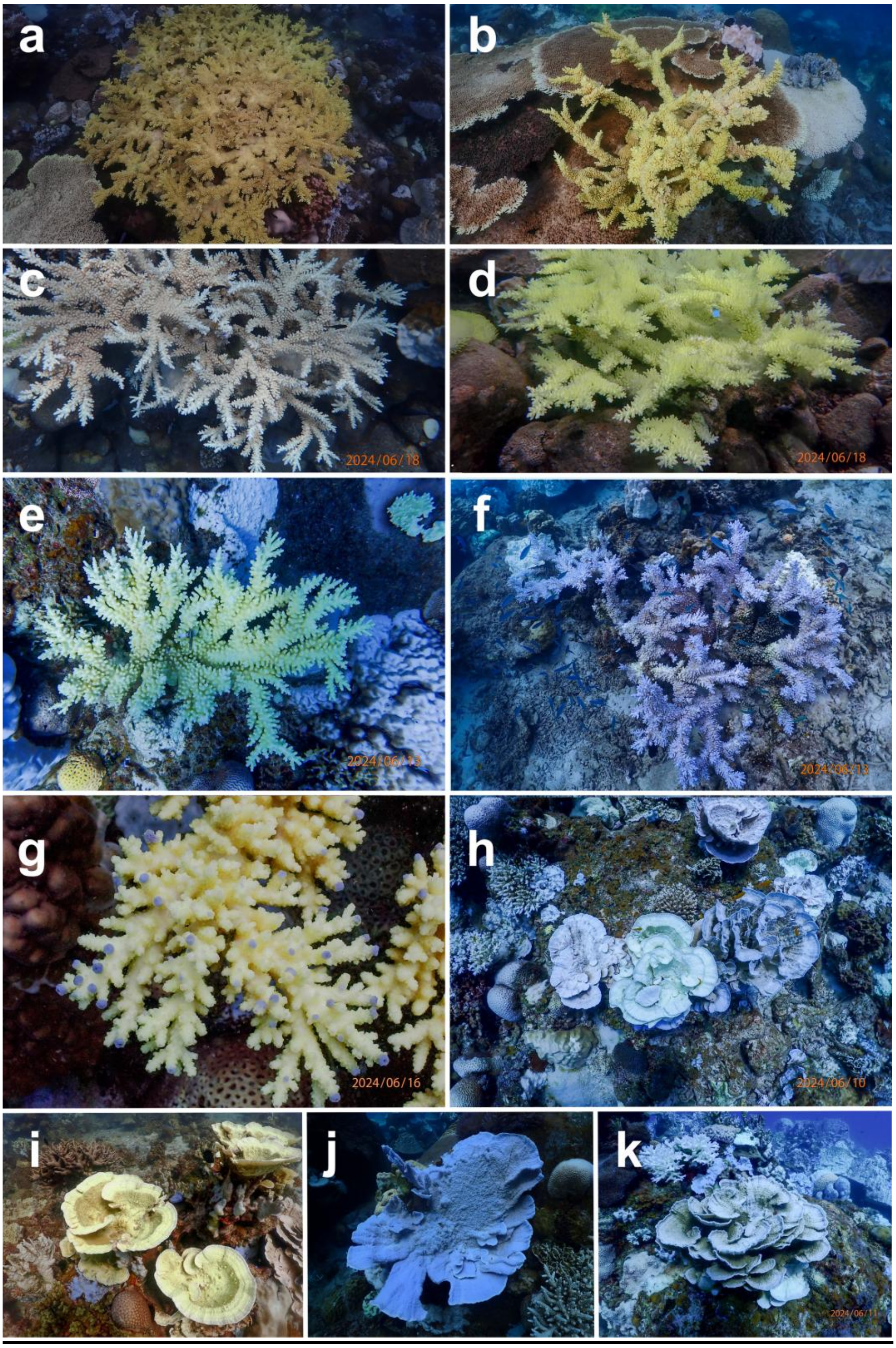


**Supplementary Figure 2. Example of coral bleaching categories.** Images (a-g) shows visual bleaching differences for *Acropora florida* colonies (photographed in June). Image (a) mild partial paling of the colonies interior; (b) proportionally higher paling of the coral colony compared to image (a). Image (c) shows severe paling and bleaching of branch ends. Image (d) shows mild colorful bleaching, compared to image (e) where the colony shows severe loss of pigmentation and bright glowing. Image (f) shows severe colourful bleaching as tissue severely depigmented and bright-blue colouration and glows is visible. Image (g) close-up photograph of severe colourful bleaching, with purple branch tips. Images (h-k) show similar variation in bleaching for *Montipora aequituberculata*

colonies. Image (h) three colonies side by side, the left-hand colony is pale, the middle is severely depigmented, and a faint yellow glow is visible (i.e., severe colourful bleaching), and the right-hand colony is glowing in blue but tissue pigmentation is predominant (i.e., mild colourful bleaching). Image (i) show a yellow to brownish glowing colony with reduced pigmentation. Image (j) a bright blue glowing colony. Image (k) partial paling and bleaching within the same colony.

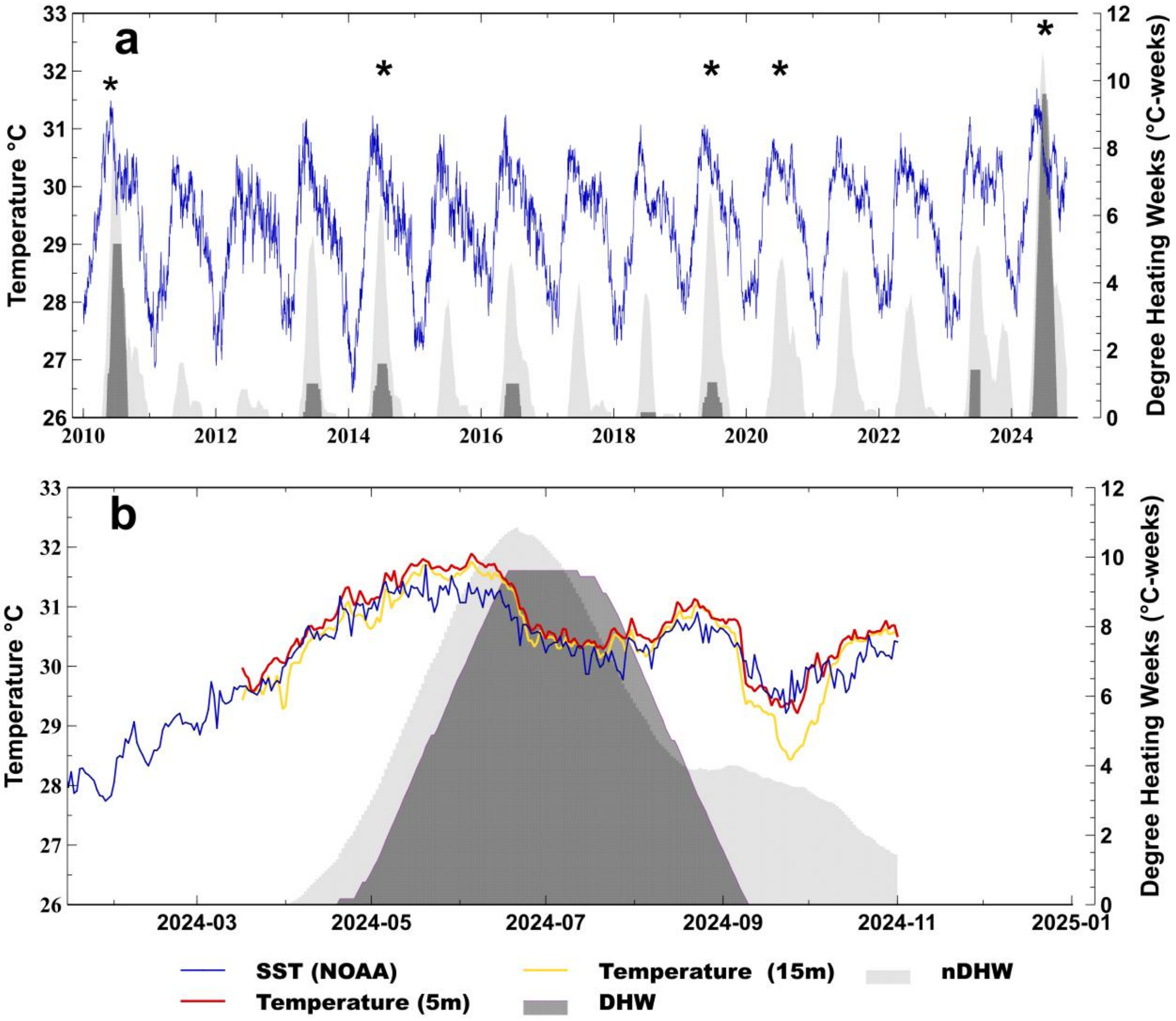


**Supplementary Figure 3. Sea surface temperature (SST) and degree heating weeks (DHW) comparison.** The blue line shows the satellite-based nightly sea surface temperatures (SST °C) around Pulau Lang Tengah, recorded by the National Oceanic and Atmospheric Association (NOAA), Coral Reef Watch (CRW) product, version 3.1. Time series are plotted against the standard DHW metric (dark-grey area), and the adjusted nDHW metric (light-grey area). Panel (a) shows heat stress events since 2010 with observed coral bleaching events (identified by available literature and author observations) highlighted by an asterisk (*). Panel (b) shows average nightly satellite-based (blue line) and in situ sea temperature measurements from five sites measured at 5 m (red line) and 15 m (yellow line) water depth, respectively, recorded between 17 March 2024, and 1 November 2024.

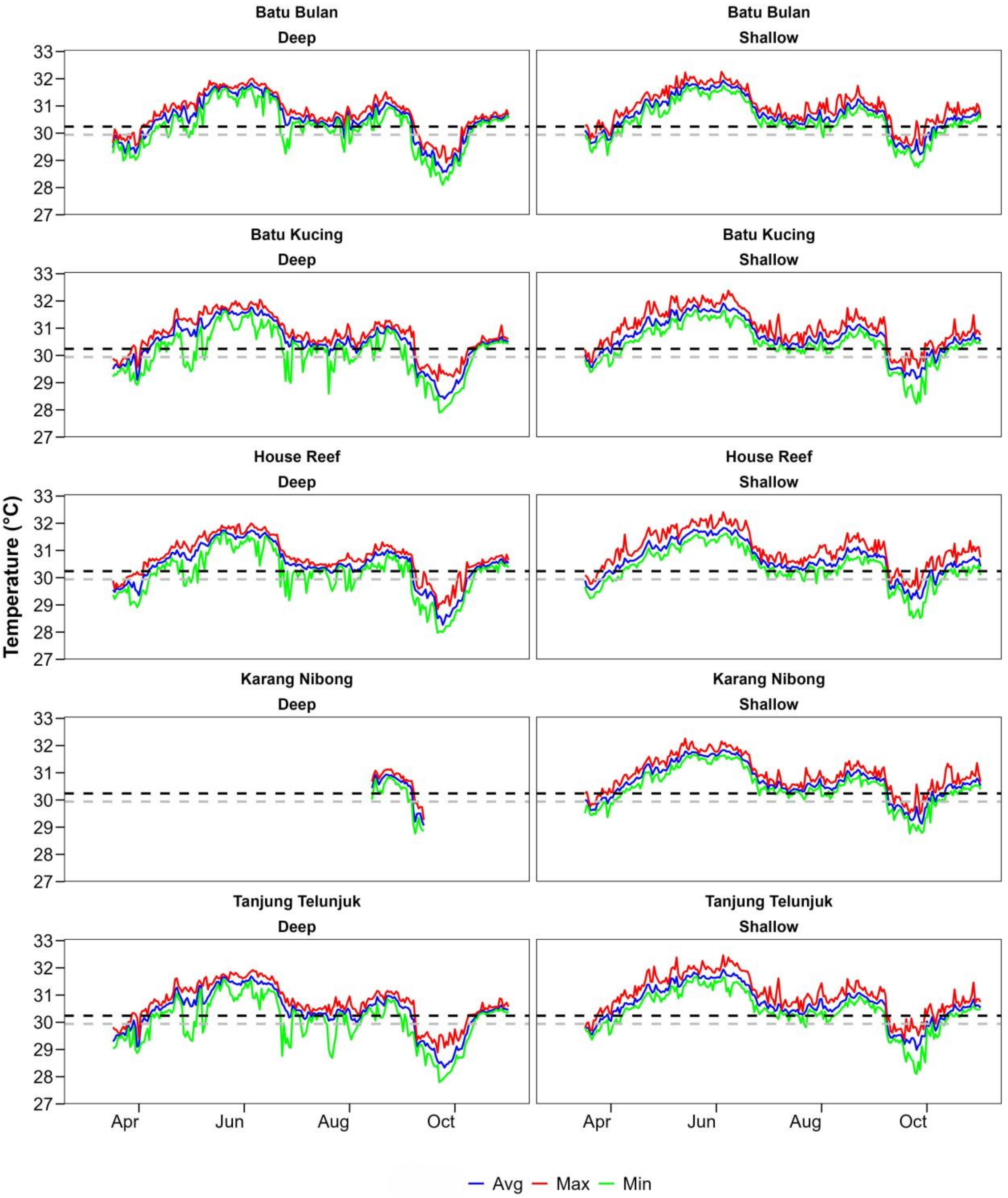


**Supplementary Figure 4. Diel temperature profiles of five sites around Pulau Lang Tengah, Malaysia.** The grey dotted line presents the maximum monthly mean (MMM=29.94°C) based on the National Oceanic and Atmospheric Association (NOAA), Coral Reef Watch (CRW) product, version 3.1, and the black dotted line presents the in-situ MMM (30.24°C) after interpolating satellite data with in-situ measurements (see Szereday et al. 2024 for methodology). The blue line shows daily mean temperature, and the green and red trend lines show daily minimum and maximum mean temperatures, respectively. Temperature data were measured at 5 m (shallow) and 15 m (deep) water depth at each site. Note, Karang Nibong deep data is missing due to logging failure.

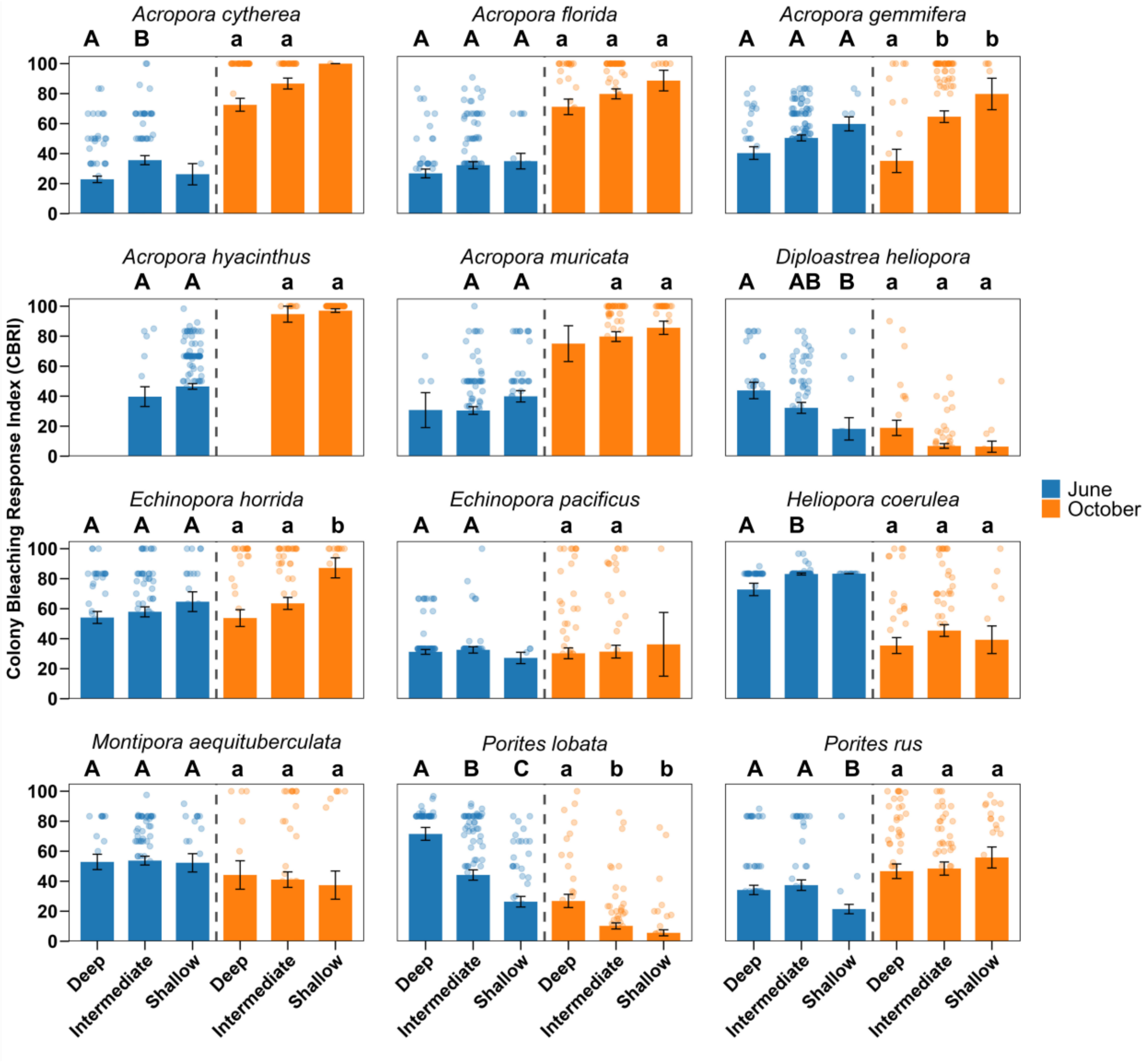


**Supplementary Figure 5. Mean differences and multiple comparisons of coral bleaching severity across depth.** Coral bleaching severity (i.e., Colony Bleaching Response Index) was measured in June 2024 during peak heat stress and four months later in October 2024. These measurements reflect immediate heat stress response and subsequent mortality, respectively. Error bars signify the standard error (±), and dots represent individual data points of coral colonies. Annotated letters alternate in each panel between upper (i.e., June) and lower case (i.e., October) letters for clarity. Letters represent results of pairwise Dunn's Tests conducted at significance levels of $p \leq 0.05$. Depth ranges are Shallow 1-5 m, Intermediate 5-10 m, Deep > 10 m.

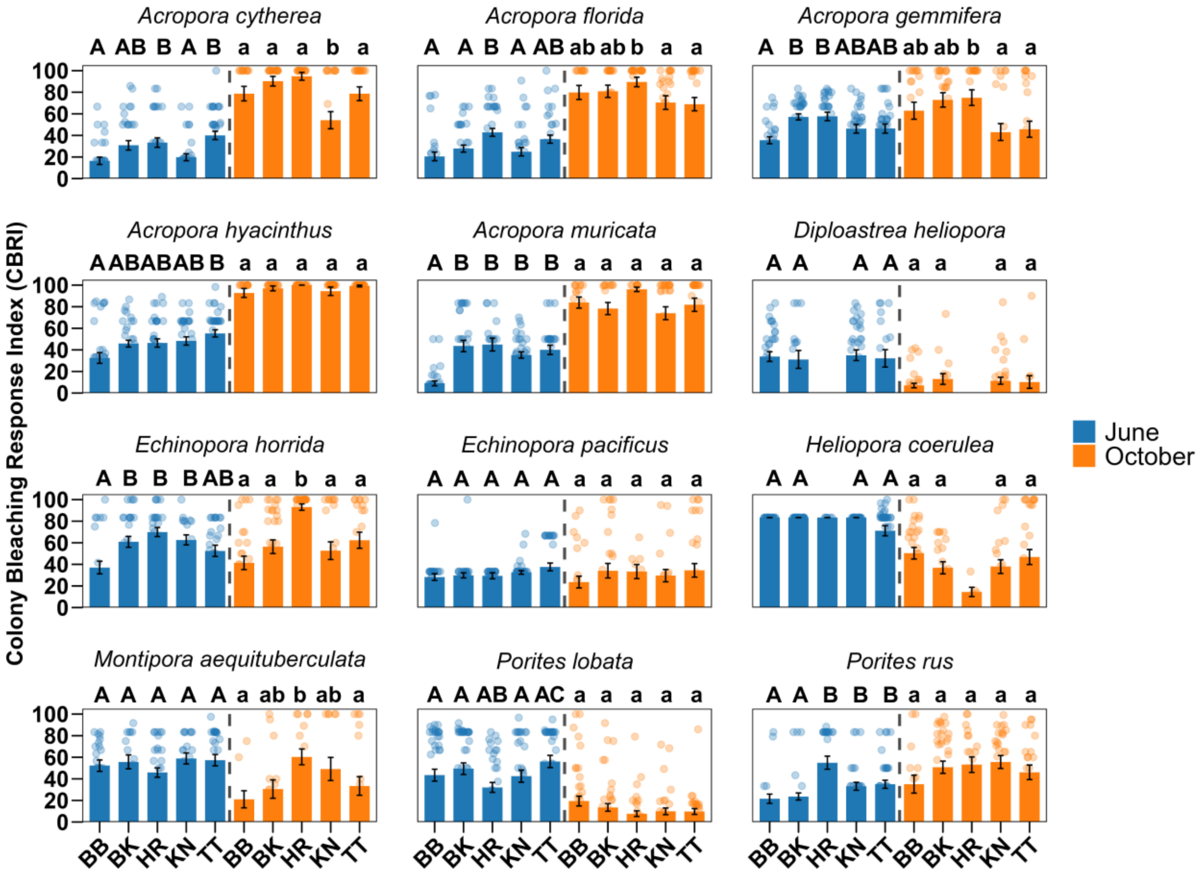


**Supplementary Figure 6. Mean differences and multiple comparisons of coral bleaching severity across sites.** Coral bleaching severity (i.e., Colony Bleaching Response Index) was measured in June 2024 during peak heat stress and four months later in October 2024. These measurements reflect immediate heat stress response and subsequent mortality, respectively. Error bars signify the standard error (±), and dots represent individual data points of coral colonies. Annotated letters alternate in each panel between upper (i.e., June) and lower case (i.e., October) letters for clarity. Letters represent results of pairwise Dunn's Tests conducted at significance levels of $p \leq 0.05$. Site abbreviations BB – Batu Bulan, BK – Batu Kucing, TT – Tanjung Telunjuk, HR – House Reef, KN – Karang Nibong

**Supplementary Table 1. Number of coral colonies surveyed per site in October 2024.** Site abbreviations BB – Batu Bulan, BK – Batu Kucing, TT – Tanjung Telunjuk, HR – House Reef, KN – Karang Nibong

| Species | BB | BK | HR | KN | TT |
|---|---|---|---|---|---|
| *Acropora cytherea* | 33 | 35 | 35 | 35 | 36 |
| *Acropora florida* | 33 | 36 | 38 | 35 | 36 |
| *Acropora gemmifera* | 30 | 34 | 34 | 32 | 34 |
| *Acropora hyacinthus* | 32 | 35 | 36 | 33 | 37 |
| *Acropora muricata* | 32 | 33 | 18 | 34 | 34 |
| *Diploastrea heliopora* | 35 | 16 | 0 | 36 | 17 |
| *Echinopora horrida* | 29 | 35 | 40 | 25 | 32 |
| *Echinopora pacificus* | 35 | 34 | 31 | 36 | 33 |
| *Heliopora coerluea* | 30 | 34 | 6 | 30 | 34 |
| *Montipora aequituberculata* | 19 | 16 | 29 | 20 | 24 |
| *Porites lobata* | 43 | 44 | 37 | 35 | 34 |

**Supplementary Table 2. Number of coral colonies surveyed per species in October 2024.** Specific depth ranges are Shallow 1-5 m, Intermediate 5-10 m, Deep > 10 m.

| Species | Shallow | Intermediate | Deep |
|---|---|---|---|
| *Acropora cytherea* | 3 | 75 | 96 |
| *Acropora florida* | 16 | 109 | 53 |
| *Acropora gemmifera* | 11 | 122 | 31 |
| *Acropora hyacinthus* | 49 | 96 | 6 |
| *Acropora muricata* | 14 | 62 | 28 |
| *Diploastrea heliopora* | 21 | 87 | 53 |
| *Echinopora horrida* | 4 | 76 | 89 |
| *Echinopora pacificus* | 13 | 78 | 43 |
| *Heliopora coerluea* | 20 | 68 | 20 |
| *Montipora aequituberculata* | 57 | 95 | 41 |
| *Porites lobata* | 24 | 57 | 60 |

**Supplementary Table 3. Multiple comparisons of heat stress across sites and depth.** Dunn's test was used to compare degree heating week (i.e., nDHW °C-Weeks) timeseries data between March 17 and November 1, 2024. Site abbreviations BB – Batu Bulan, BK – Batu Kucing, TT – Tanjung Telunjuk, HR – House Reef, KN – Karang Nibong. Specific depth ranges are Shallow 1-5 m, Intermediate 5-10 m, Deep > 10 m. SE- standard error.

| *Site 1* | *Site 2* | *R-mean* | *SE* | *z-stat* | *R-crit* | *p-value* |
|---|---|---|---|---|---|---|
| BB Shallow | BB Deep | 181.0196 | 55.71999 | 3.248736 | 109.2092 | 0.001159 |
| BB Shallow | BK Shallow | 2.252174 | 55.71999 | 0.040419 | 109.2092 | 0.967759 |
| BB Shallow | BK Deep | 198.3174 | 55.71999 | 3.559178 | 109.2092 | 0.000372 |
| BB Shallow | HR Shallow | 46.94783 | 55.71999 | 0.842567 | 109.2092 | 0.399471 |
| BB Shallow | HR Deep | 197.1783 | 55.71999 | 3.538734 | 109.2092 | 0.000402 |
| BB Shallow | KN Shallow | 33.96522 | 55.71999 | 0.60957 | 109.2092 | 0.542147 |
| BB Shallow | TT Shallow | 21.95217 | 55.71999 | 0.393973 | 109.2092 | 0.693601 |
| BB Shallow | TT Deep | 318.0913 | 55.71999 | 5.708746 | 109.2092 | 1.14E-08 |
| BB Deep | BK Shallow | 178.7674 | 55.71999 | 3.208317 | 109.2092 | 0.001335 |
| BB Deep | BK Deep | 17.29783 | 55.71999 | 0.310442 | 109.2092 | 0.756225 |
| BB Deep | HR Shallow | 134.0717 | 55.71999 | 2.406169 | 109.2092 | 0.016121 |
| BB Deep | HR Deep | 16.1587 | 55.71999 | 0.289998 | 109.2092 | 0.771818 |
| BB Deep | KN Shallow | 147.0543 | 55.71999 | 2.639167 | 109.2092 | 0.008311 |
| BB Deep | TT Shallow | 159.0674 | 55.71999 | 2.854763 | 109.2092 | 0.004307 |
| BB Deep | TT Deep | 137.0717 | 55.71999 | 2.46001 | 109.2092 | 0.013893 |
| BK Shallow | BK Deep | 196.0652 | 55.71999 | 3.518759 | 109.2092 | 0.000434 |
| BK Shallow | HR Shallow | 44.69565 | 55.71999 | 0.802147 | 109.2092 | 0.422468 |
| BK Shallow | HR Deep | 194.9261 | 55.71999 | 3.498315 | 109.2092 | 0.000468 |
| BK Shallow | KN Shallow | 31.71304 | 55.71999 | 0.56915 | 109.2092 | 0.569254 |
| BK Shallow | TT Shallow | 19.7 | 55.71999 | 0.353554 | 109.2092 | 0.723674 |
| BK Shallow | TT Deep | 315.8391 | 55.71999 | 5.668327 | 109.2092 | 1.44E-08 |
| BK Deep | HR Shallow | 151.3696 | 55.71999 | 2.716611 | 109.2092 | 0.006595 |
| BK Deep | HR Deep | 1.13913 | 55.71999 | 0.020444 | 109.2092 | 0.983689 |
| BK Deep | KN Shallow | 164.3522 | 55.71999 | 2.949609 | 109.2092 | 0.003182 |
| BK Deep | TT Shallow | 176.3652 | 55.71999 | 3.165205 | 109.2092 | 0.00155 |
| BK Deep | TT Deep | 119.7739 | 55.71999 | 2.149568 | 109.2092 | 0.031589 |
| HR Shallow | HR Deep | 150.2304 | 55.71999 | 2.696167 | 109.2092 | 0.007014 |
| HR Shallow | KN Shallow | 12.98261 | 55.71999 | 0.232997 | 109.2092 | 0.815763 |
| HR Shallow | TT Shallow | 24.99565 | 55.71999 | 0.448594 | 109.2092 | 0.653725 |
| HR Shallow | TT Deep | 271.1435 | 55.71999 | 4.866179 | 109.2092 | 1.14E-06 |
| HR Deep | KN Shallow | 163.213 | 55.71999 | 2.929165 | 109.2092 | 0.003399 |
| HR Deep | TT Shallow | 175.2261 | 55.71999 | 3.144761 | 109.2092 | 0.001662 |
| HR Deep | TT Deep | 120.913 | 55.71999 | 2.170012 | 109.2092 | 0.030006 |
| KN Shallow | TT Shallow | 12.01304 | 55.71999 | 0.215597 | 109.2092 | 0.829302 |
| KN Shallow | TT Deep | 284.1261 | 55.71999 | 5.099177 | 109.2092 | 3.41E-07 |
| TT Shallow | TT Deep | 296.1391 | 55.71999 | 5.314773 | 109.2092 | 1.07E-07 |

**Supplementary Table 4. Kruskal-Wallis test of species-specific depth effects on coral bleaching outcomes in October 2024.** Specific depth ranges are Shallow 1-5 m, Intermediate 5-10 m, Deep > 10 m. Sample size per depth range is shown. KW - Kruskal-Wallis; df – degree of freedom.

| Species | Shallow | Intermediate | Deep | KW | df | p-value |
|---|---|---|---|---|---|---|
| *Acropora cytherea* | 3 | 75 | 96 | 5.96 | 2 | 0.0508 |
| *Acropora florida* | 16 | 109 | 53 | 4.39 | 2 | 0.1113 |
| *Acropora gemmifera* | 11 | 122 | 31 | 14.48 | 2 | 7.00E-04 |
| *Acropora hyacinthus* | 49 | 96 | 6 | 3.37 | 2 | 0.1851 |
| *Acropora muricata* | 14 | 62 | 28 | 3.82 | 2 | 0.1484 |
| *Diploastrea heliopora* | 21 | 87 | 53 | 11.92 | 2 | 0.0026 |
| *Echinopora horrida* | 4 | 76 | 89 | 1.21 | 2 | 0.5465 |
| *Echinopora pacificus* | 13 | 78 | 43 | 3.77 | 2 | 0.1515 |
| *Heliopora coerluea* | 20 | 68 | 20 | 0.84 | 2 | 0.6561 |
| *Montipora aequituberculata* | 57 | 95 | 41 | 36.00 | 2 | 0 |
| *Porites lobata* | 24 | 57 | 60 | 1.38 | 2 | 0.5029 |

**Supplementary Table 5. Kruskal-Wallis test of species-specific site effects on coral bleaching outcomes in October 2024.** Sample size per depth range is shown. Site abbreviations BB – Batu Bulan, BK – Batu Kucing, TT – Tanjung Telunjuk, HR – House Reef, KN – Karang Nibong KW - Kruskal-Wallis; df – degree of freedom.

| Species | BB | BK | HR | KN | TT | KW | df | p-value |
|---|---|---|---|---|---|---|---|---|
| *Acropora cytherea* | 33 | 35 | 35 | 35 | 36 | 23.47 | 4 | 1.00E-04 |
| *Acropora florida* | 33 | 36 | 38 | 35 | 36 | 16.15 | 4 | 0.0028 |
| *Acropora gemmifera* | 30 | 34 | 34 | 32 | 34 | 15.89 | 4 | 0.0032 |
| *Acropora hyacinthus* | 32 | 35 | 36 | 33 | 37 | 4.02 | 4 | 0.4032 |
| *Acropora muricata* | 32 | 33 | 18 | 34 | 34 | 6.65 | 4 | 0.1555 |
| *Diploastrea heliopora* | 35 | 16 | 0 | 36 | 17 | 2.21 | 3 | 0.5307 |
| *Echinopora horrida* | 29 | 35 | 40 | 25 | 32 | 41.95 | 4 | 0 |
| *Echinopora pacificus* | 35 | 34 | 31 | 36 | 33 | 5.42 | 4 | 0.2472 |
| *Heliopora coerluea* | 30 | 34 | 6 | 30 | 34 | 7.79 | 4 | 0.0998 |
| *Montipora aequituberculata* | 19 | 16 | 29 | 20 | 24 | 16.73 | 4 | 0.0022 |
| *Porites lobata* | 43 | 44 | 37 | 35 | 34 | 3.056 | 4 | 0.5485 |
